# Enhanced SARS-CoV-2 Neutralization by Secretory IgA in vitro

**DOI:** 10.1101/2020.09.09.288555

**Authors:** Zijun Wang, Julio C. C. Lorenzi, Frauke Muecksch, Shlomo Finkin, Charlotte Viant, Christian Gaebler, Melissa Cipolla, Hans-Heinrich Hoffman, Thiago Y. Oliveira, Deena A. Oren, Victor Ramos, Lilian Nogueira, Eleftherios Michailidis, Davide F. Robbiani, Anna Gazumyan, Charles M. Rice, Theodora Hatziioannou, Paul D. Bieniasz, Marina Caskey, Michel C. Nussenzweig

## Abstract

SARS-CoV-2 primarily infects cells at mucosal surfaces. Serum neutralizing antibody responses are variable and generally low in individuals that suffer mild forms of the illness. Although potent IgG antibodies can neutralize the virus, less is known about secretory antibodies such as IgA that might impact the initial viral spread and transmissibility from the mucosa. Here we characterize the IgA response to SARS-CoV-2 in a cohort of 149 individuals. IgA responses in plasma generally correlate with IgG responses and clones of IgM, IgG and IgA producing B cells that are derived from common progenitors are evident. Plasma IgA monomers are 2-fold less potent than IgG equivalents. However, IgA dimers, the primary form in the nasopharynx, are on average 15 times more potent than IgA monomers. Thus, secretory IgA responses may be particularly valuable for protection against SARS-CoV-2 and for vaccine efficacy.

## Introduction

SARS-CoV-2 encodes a trimeric spike surface protein (S) which mediates entry into host cells (*1, 2*). The virus initially infects epithelial cells in the nasopharynx when the receptor binding domain (RBD) of S interacts with angiotensin converting enzyme-2 (ACE-2) receptor (*3-6*). SARS-CoV-2 may subsequently spread to other epithelial cells expressing ACE-2 in the lung and gut. These tissues are rich in lymphoid cells that are organized into nasopharynx associated and gut associated lymphoid tissues (NALT and GALT respectively). Vaccines delivered by inhalation to specifically target these tissues appear to be more effective against SARS-CoV-2 (*7*). Among other specializations, NALT and GALT produce large quantities of IgA antibodies. These antibodies exist as monomers in circulation where they make up 15% of the serum antibody pool. However, IgA is found in higher concentrations in secretions where it exists predominantly as a dimer covalently linked by J chain (*8-10*).

Although most individuals produce antibodies in response to SARS-CoV-2 infection, the neutralizing response is highly variable with as many as 30% of the population showing levels of neutralizing activity below 1:50 in pseudovirus assays (*11, 12*). Neutralization is associated with prolonged infection and RBD binding activity as measured by ELISA (*11-13*). IgG antibody cloning experiments from recovered individuals have revealed that neutralizing antibodies target several distinct non-overlapping epitopes on the RBD (*11, 14-18*). Some of these antibodies are potently neutralizing and can prevent or treat infection in animal models (*15-19*).

Consistent with the fact that SARS CoV-2 initially infects in the nasopharynx, IgA antibodies that bind to SARS-CoV-2 are produced rapidly after infection and remain elevated in the plasma for at least 40 days after the onset of symptoms (*20-23*). IgA antibodies bind to the RBD and can neutralize SARS-CoV-2 (*20-22*). However, the precise contribution and molecular nature of the IgA response to SARS-CoV-2 has not been reported to date. Here we examine a cohort of 149 convalescent individuals with measurable plasma neutralizing activity for the contribution of IgA to anti-SARS-CoV-2 antibody responses. Cloning IgA antibodies from single B cells reveals that the neutralizing activity of monomeric IgA is generally lower than corresponding IgGs but dimeric IgAs are on average 15-fold more potent than their monomeric counterparts.

## Results

### Plasma anti-SARS-CoV-2 RBD IgA

IgM, IgG and IgA account for 5%, 80% and 15% of the antibodies in plasma, respectively. IgG responses to RBD are strongly correlated with neutralizing activity (*11, 13-17, 24-28*). To examine the contribution of IgA to the anti-SARS-CoV-2 RBD response we tested plasma samples for binding to the RBD by a validated ELISA. A positive control sample (COV-21) was included for normalization of the area under the curve (AUC) and 8 independent healthy donor samples were included as negative controls (Fig. 1A, (*11*)). Whereas 78% and 15% of the individuals in this cohort showed IgG and IgM anti-RBD levels that were at least 2 standard deviations above control, only 33% did so for IgA (Fig. 1A and B, (*11*)). Thus, in individuals studied on average 40 days after infection the circulating levels of anti-RBD IgA is more modest than IgG and higher than IgM.

**Fig. 1.**
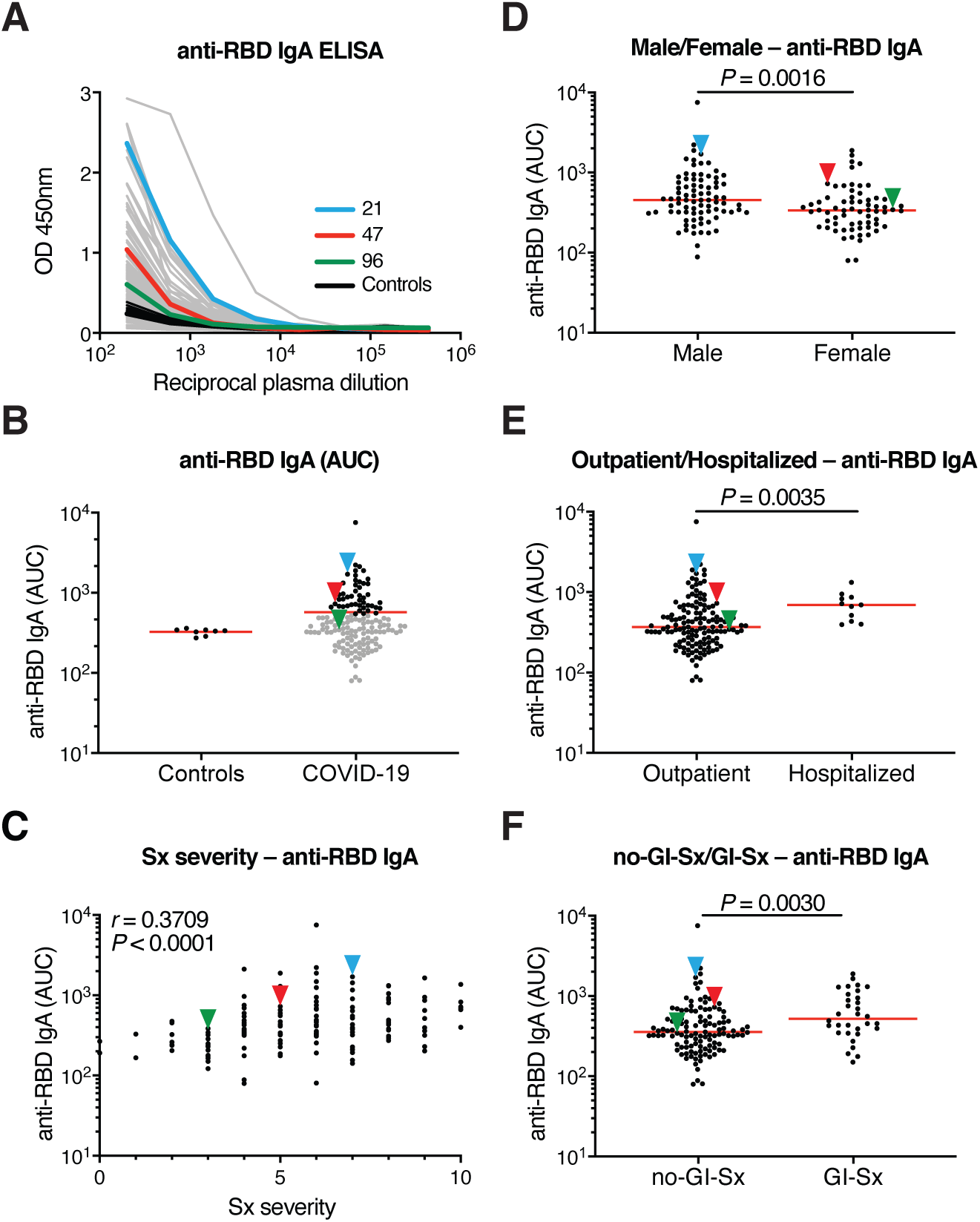
Plasma IgA against SARS-CoV-2 RBD. (A) ELISAs measuring plasma IgA reactivity to RBD. Graph shows optical density units at 450 nm (OD, Y axis) and reciprocal plasma dilutions (X axis). Negative controls in black; individuals 21, 47, 96 in blue, red and green lines and arrowheads, respectively (*11*). (B) Graph shows normalized area under the curve (AUC) for 8 controls and each of 149 individuals in the cohort. Horizontal bar indicates mean values. Black dots indicate the individuals that are 2 STDV over the mean of controls. (C) Subjective Symptom (Sx) severity (X axis) is plotted against the normalized AUC for IgA binding to RBD (Y axis). *R* = 0.3709, *P* < 0.0001. (D) Normalized AUC of anti-RBD IgA ELISA for males (n=83) and females (n=66); *P* =0.0016. (E) Normalized AUC of anti-RBD IgA ELISA for outpatients (n=138) and hospitalized (n=11) individuals; *P* = 0.0035. (F) Normalized AUC of anti-RBD IgA ELISA for patients with gastrointestinal (GI) symptoms (n=32) and without GI symptoms (n=117); *P* = 0.0030. The *r* and *P* values for the correlations in (C) were determined by two-tailed Spearman’s. For (D-F) horizontal bars indicate median values. Statistical significance was determined using two-tailed Mann-Whitney U test.

Anti-RBD IgA titers were correlated with duration and severity of symptoms but not timing of sample collection relative to onset (Fig. 1C, and fig.S1A, B). Similar to IgG, females had lower levels of IgA than males and hospitalized individuals showed higher anti-RBD IgA titers than those with milder symptoms, but there was no correlation with age (Fig. 1D and E, fig. S1C). Of note, individuals that suffered gastrointestinal symptoms showed significantly higher plasma anti-RBD IgA but not IgG titers (Fig. 1F and fig. S1D).

### Neutralization activity of purified IgG and IgA

To compare the neutralizing activity of plasma IgA to IgG directly we purified the 2 isotypes from the plasma of all 99 individuals in our cohort that showed measurable plasma neutralizing activity and tested the two isotypes in HIV-1 based SARS-CoV-2 pseudovirus neutralization assays (*11*). The activity of both isotypes was directly correlated with anti-RBD binding titers and overall plasma neutralizing activity (Fig. 2A-D). In addition, there was good correlation between the neutralizing activity of IgG and IgA in a given individual (Fig. 2E). However, potency of each of the 2 isotypes varied by as much as 2 orders of magnitude between individuals (Fig. 2F). Purified IgG was generally more potent than IgA in neutralizing SARS-CoV-2 pseudovirus *in vitro*. The geometric mean IC_50_ for IgG was 384 nM vs. 709 nM for IgA (*P* < 0.0001, Fig. 2F). Nevertheless, IgAs were more potent than IgGs in 25% of the individuals tested (fig. S2A). The 2 isotypes also differed in that the overall potency of purified IgG was correlated with symptom severity and was higher in hospitalized individuals, but purified IgA was not (Fig. 2G and fig. S2B-D). Finally, the potency of the purified IgA was greater in individuals that suffered from gastrointestinal symptoms, but IgG was not (Fig. 2H and fig. S2E).

**Fig. 2.**
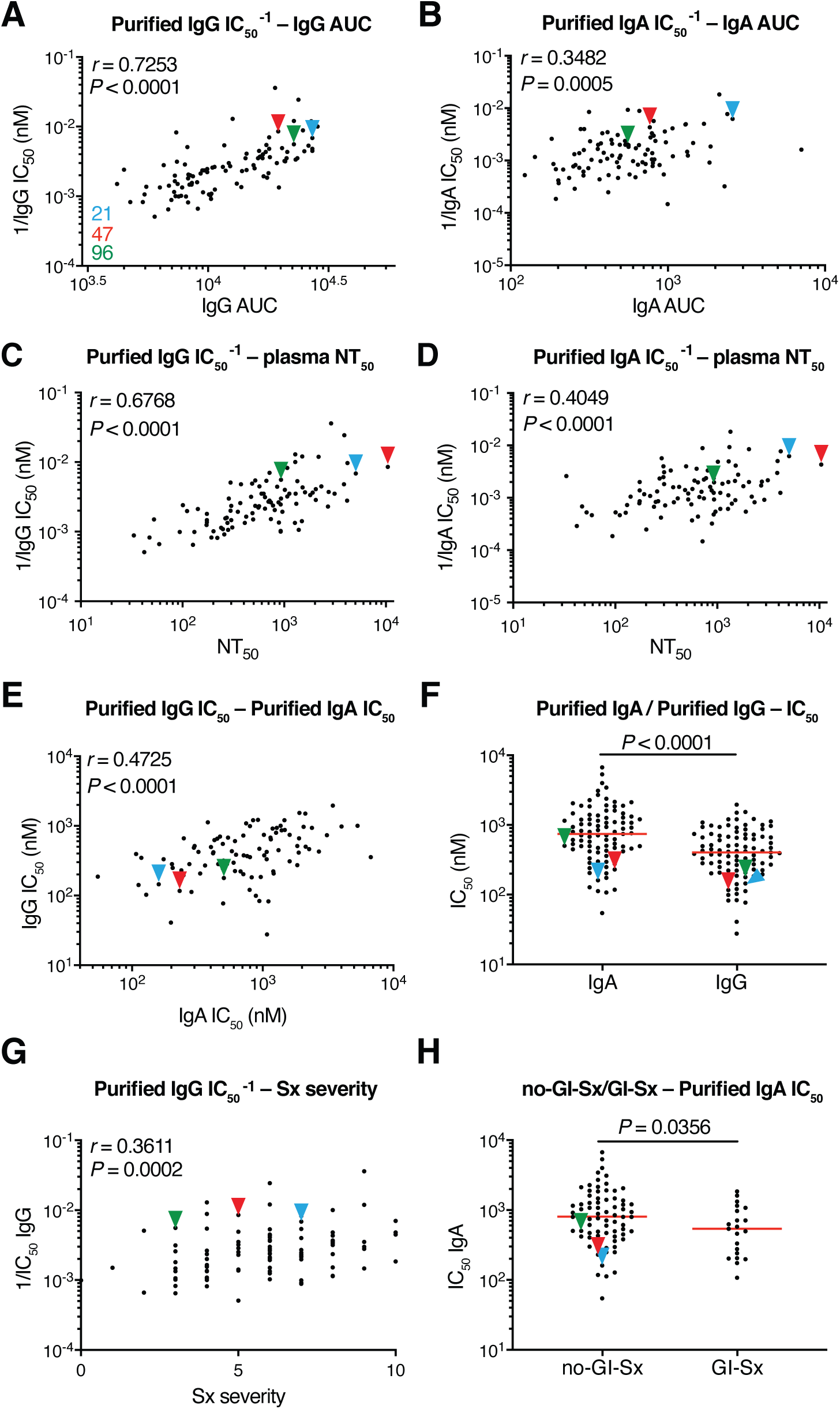
SARS-CoV-2 pseudovirus neutralization by purified IgA and IgG. Neutralization activity of plasma-purified IgG and IgA from 99 participants measured in cell lysates of HT1080_ACE2_cl.14 cells 48 h after infection with nanoluc-expressing SARS-CoV-2 pseudovirus. (A) Normalized AUC for plasma IgG anti-RBD ELISA (X axis) plotted against purified IgG pseudovirus neutralization 1/IC_50_ values (Y axis). *r* = 0.7253, *P* < 0.0001. (B) Normalized AUC for plasma IgA ELISA (X axis) plotted against purified IgA pseudovirus neutralization 1/IC_50_ values (Y axis). *r* = 0.3482, *P* = 0.0005. (C) Published plasma NT_50_ values (*11*) (X axis) plotted against purified IgG pseudovirus neutralization 1/IC_50_ values (Y axis). *r* = 0.6768, *P* < 0.0001. (D) Published plasma NT_50_ values (*11*) (X axis) plotted against purified IgA pseudovirus neutralization 1/IC_50_ values (Y axis). *r* = 0.4049, *P* < 0.0001. (E) Purified IgA pseudovirus neutralization IC_50_ values (X axis) plotted against purified IgG pseudovirus neutralization IC_50_ values. *r* = 0.4725, *P* < 0.0001. (F) Comparison of purified IgA and IgG pseudovirus neutralization IC_50_ values, *P* < 0.0001. (G) Symptom severity plotted against purified IgG pseudovirus neutralization 1/IC_50_ values. *r* = 0.3611, *P* = 0.0002. (H) Purified IgA pseudovirus neutralization IC_50_ values for patients with GI symptoms (n=21) and without GI symptoms (n=74); *P* = 0.0356. The *r* and *p* values in (A-E, G) were determined by two-tailed Spearman’s correlations. In (F and H), *p* values were determined by two-tailed Mann–Whitney U-tests and horizontal bars indicate median values.

### Monoclonal anti-SARS-CoV-2 IgM and IgA antibodies

To determine the nature of the IgM and IgA anti-RBD antibodies elicited by SARS-CoV-2 infection we used flow cytometry to purify single B lymphocytes that bind to RBD and cloned their antibodies. We obtained 109 IgM and 74 IgA (64 IgA1 and 10 IgA2) matched Ig heavy and light chain sequences by reverse transcription and subsequent isotype specific PCR from 3 convalescent individuals (Fig. 3A, B). As reported for IgG antibodies (*11, 14, 17, 26, 29*), the overall number of mutations was generally low when compared to antibodies obtained from individuals suffering from chronic infections such as Hepatitis-B or HIV-1 (*30, 31*) (fig. S3A, B). However, the number of V gene nucleotide mutations in IgM and IgA heavy and light chains varied between individuals. For example, in donor COV21 the number of IgM and IgA heavy chain mutations was similar. In contrast, IgM heavy and light chain nucleotide mutations were significantly greater than IgA mutations in COV47 (fig. S3B). CDR3 length was significantly shorter for IgM than IgA and IgG antibodies and hydrophobicity was slightly higher for IgM over control but not for IgA and IgG (figs. S4 and S5). Compared to the normal human antibody repertoire, several IgA and IgM VH genes were over-represented including VH3-53 which can make key contacts with the RBD through germline encoded CDRH1 and CDRH2 (*11, 32, 33*) (fig. S6).

**Fig. 3.**
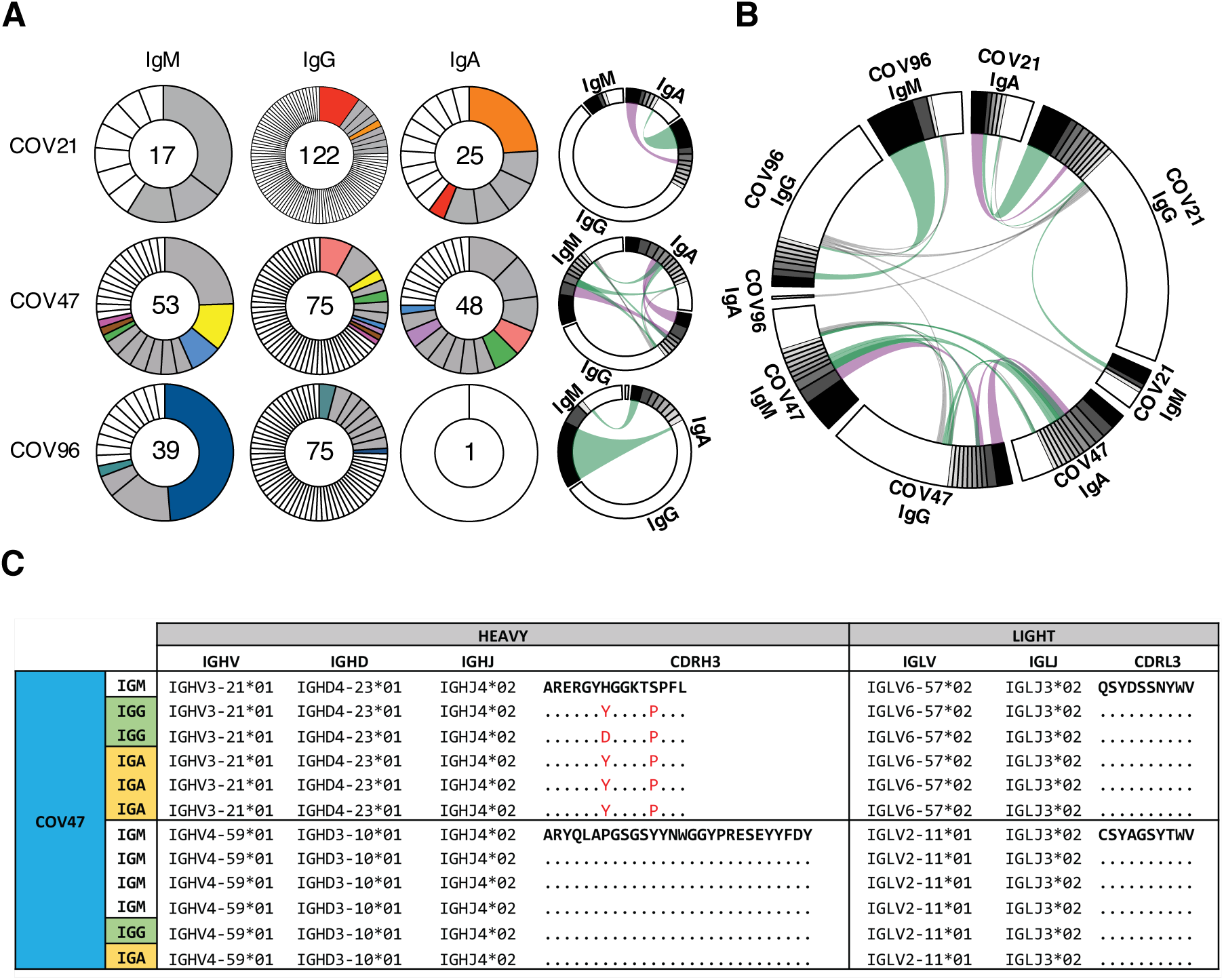
Monoclonal anti-SARS-CoV-2 RBD IgM, IgG and IgA. (A) Clonal expansion of B cells producing of IgM, IgG and IgA from three individuals. The number in the inner circle indicates the number of sequences analyzed for the individual denoted above the circle. Pie slices size is proportional to the number of clonally related sequences. Colored pie slices indicate clones or singlets that share the same IGHV and IGLV genes, and highly similar CDR3s. Grey indicates clones that are not shared. White indicates singlets that are not shared. The right side circos plots show the relationship between antibodies of different isotypes that share same IGH V(D)J and IGL VJ genes, and highly similar CDR3s. Purple, green and grey lines connect related clones, clones and singles, and singles to each other, respectively. (B) Circos plot shows sequences from all 3 individuals with clonal relationships depicted as in (A). (C) Sample sequence alignment for antibodies of different isotypes that display same IGH V(D)J and IGL VJ genes and highly similar CDR3s. Amino acid differences in CDR3s to the reference sequence (bold) are indicated in red, dashes indicate missing amino acids and dots represent identical amino acids.

Like IgG antibodies (*11*) IgA and IgM antibodies were found in expanded clones in all 3 of the individuals examined. Overall 66.2% and 66.1% of all the IgA and IgM sequences examined were members of expanded clones (Fig. 3A, B and table S1). Nearly identical sequences were shared among the 3 isotypes in clones found in all 3 individuals indicating that switch recombination occurred during B cell clonal expansion in response to SARS-CoV-2 (Fig.3B). In total 11 out of 55 antigen-specific B cell clones in circulation belonged to expanded clones that contained members expressing different constant regions (Fig. 3C and tables S1 and S2). When compared directly, the neutralizing activity of antibodies that were members of B cell clones producing IgA or IgG varied and did not correlate with one or the other isotype (table S3).

To examine the binding properties of the anti-SARS-CoV-2 monoclonals we expressed 46 IgMs and 35 IgAs (table S4). IgM variable regions were produced on an IgG1 backbone to facilitate expression and purification. IgAs were expressed as native IgA1 or IgA2 monomers. ELISA assays on RBD showed that 100% and 91.3% of the IgA and IgM antibodies bound to the RBD with an average half-maximal effective concentration of 52.8 ng/ml and 101.6 ng/ml respectively (fig. S7A and table S5).

To determine neutralizing activity of the IgM and IgA antibodies we tested them against an HIV-1 based SARS-CoV-2 pseudovirus as IgGs and native IgA monomers respectively. Among the 42 RBD binding IgM antibodies tested we found 10 that neutralized the virus in the ng/ml range with geometric mean half-maximal inhibitory concentrations (IC_50_) of 114.0 nanograms per milliliter (Fig. 4A and fig. S7B, table S5). In contrast, 32 out of 35 RBD binding IgA antibodies tested neutralized the virus in the ng/ml range with geometric mean half-maximal inhibitory concentrations (IC_50_) of 53.6 nanograms per milliliter (Fig. 4A and fig. S7B, table S5). Thus, IgM antibodies expressed as monomeric IgGs show lower neutralizing activity than either native IgA or IgG monomers (Fig. 4A).

**Fig. 4.**
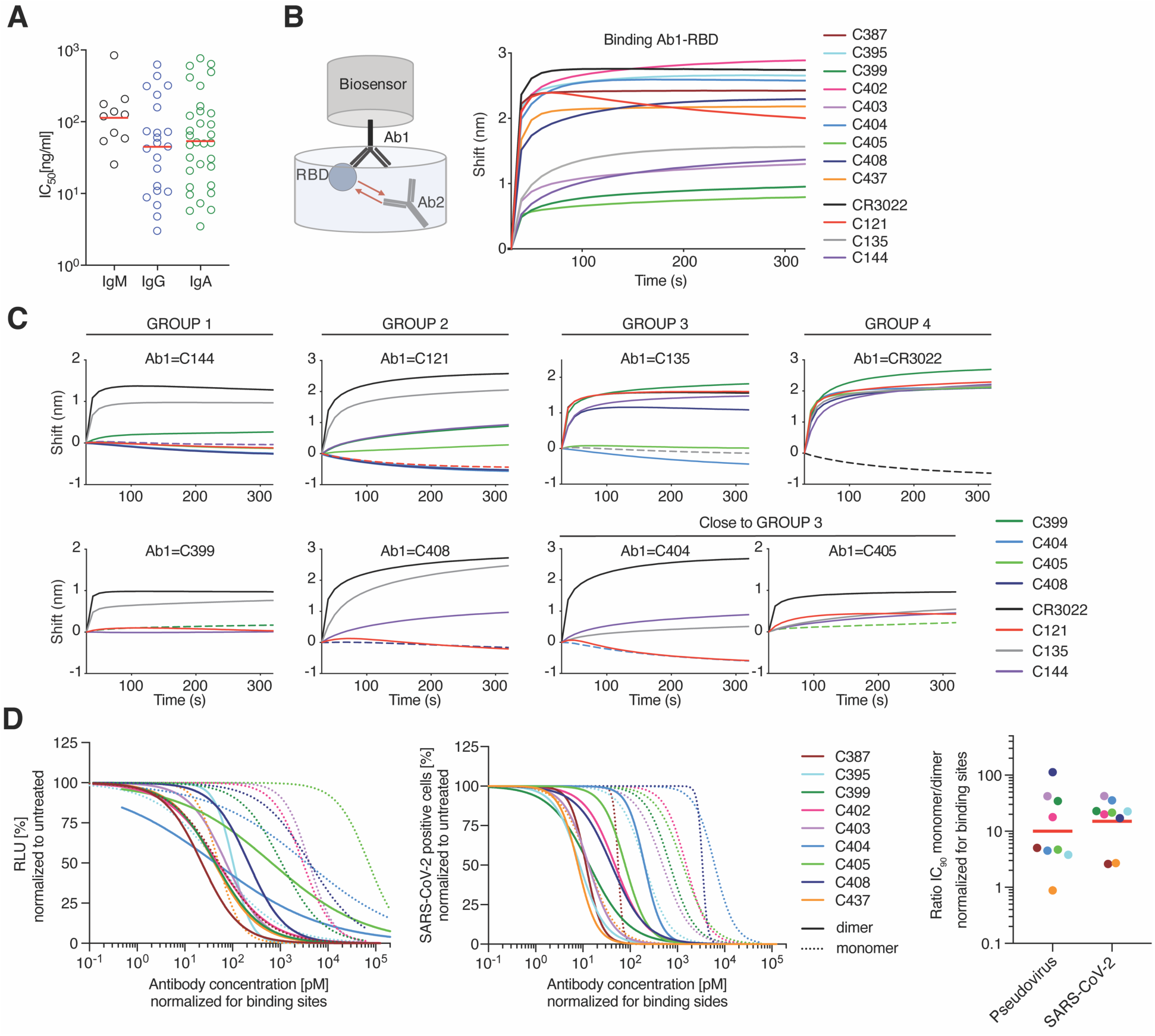
IgA dimers neutralize SARS-CoV-2 more potently than monomers. (A) Pseudovirus IC_50_ neutralization values for IgA, and IgM monoclonals and published IgG monoclonals from the same individuals (*11*). Antibodies with IC_50_ less than 1000 ng/ml are shown. Red lines indicate geometric mean. (B) Diagrammatic representation of biolayer interferometry experiment (left panel). Binding of C387, C395, C399, C402, C403, C404, C405, C408, C437, CR3022, C121, C135, C144 to RBD (right panel). (C) Second antibody (Ab2) binding to preformed first antibody (Ab1)–RBD complexes. Dotted line denotes when Ab1 and Ab2 are the same, and Ab2 is according to the colour-coding in g. h, l, Group 1 antibodies were tested. (D) The normalized relative luminescence values for cell lysates of 293T_ACE2_ cells after infection with SARS-CoV-2 pseudovirus (left panel) or normalized percentage of SARS-CoV-2 positive VeroE6 cells 48 h after infection with SARS-CoV-2 authentic virus (middle panel; values obtained in the absence of antibody are plotted at x=0.1 to be visible on log-scale) in the presence of increasing concentrations of monoclonal antibodies C387, C395, C399, C402, C403, C404, C405, C408, C437 as monomers or dimers. Shown are four-parameter nonlinear regression curve fits of normalized data. Comparison of the ratio of IC_90_ values of monomer to dimers, normalized to number of antibody binding sites (right panel).

### Dimeric anti-SARS-CoV-2 IgA is more potent than monomeric IgA

To determine whether these IgAs targeted the same epitopes as previously characterized IgGs we performed bilayer interferometry experiments in which a preformed antibody-RBD complex consisting of C144-RBD, or C121-RBD or C135-RBD or CR3022-RBD (Fig. 4B) was exposed to a monomeric IgA monoclonal. C144 and C121 recognize the ACE-2 interaction domain of the RBD, C135 and CR3022 neutralize without interfering with ACE-2 binding (Fig. 4C) (*11, 32, 34*). Two of the IgA’s were in the C144 category, 5 were similar to C121, and 2 resembled C135 (Fig. 4C and fig. S9). Thus, RBD recognition by neutralizing IgA is similar to IgG.

Mucosal IgA exists predominantly as a dimer. To examine the neutralizing activity of IgA dimers we co-expressed 8 IgA1s and 1 IgA2 with J chain to produce mixtures of monomers and dimers that were purified by size exclusion chromatography (fig. S8). When compared in pseudovirus neutralization assays, 8 out of 9 IgA dimers were more potent than the corresponding monomers with differences in activity ranging from 3.8 to 113-fold (Fig. 4D, fig. S10A and table S6). The relative increase in neutralizing activity between monomer and dimer was inversely correlated with the neutralizing activity of the monomer in this assay (fig. S10B. IC_50_: *r*=0.80, *P*=0.014). For example, whereas C437, the most potent antibody, showed equivalent activity as a monomer and dimer, C408, one of the least potent antibodies, was 113-fold more potent as a dimer (fig. S10B).

IgA monomers and dimers were also compared in SARS-CoV-2 microneutralization assays. Neutralizing activities of the 9 monomers and 9 dimers correlated strongly with those measured in the pseudovirus neutralization assay (fig. S10C. IC_50_: *r*=0.84, *P*<0.0001; IC_90_: *r*=0.91, *P*<0.0001). On average, there was a 15-fold geometric mean increase in activity for the dimer over the monomer against SARS-CoV-2 and less variability in the degree of enhancement in microneutralization compared to pseudovirus assays (Fig. 4D, fig. S10D and E, and table S6). Thus, dimeric IgA is far more potent than monomeric IgA against SARS-CoV-2 (Fig. 4D).

## Discussion

Neutralizing antibody titers are the best correlates of protection in most vaccines (*35*). Among antibody isotypes, secretory IgA which is found at mucosal surfaces, plays a crucial role in protecting against pathogens that target these surfaces (*36*). We find that serum IgA responses to SARS-CoV-2 correlate with IgG responses. Although the monomeric form of IgA found in serum is on average 2-fold less potent than IgG, the dimeric secretory form of IgA found in mucosa is over one log more potent than the monomer against authentic SARS-CoV-2 which makes it a far more potent neutralizer than IgG.

The increased potency of the dimeric form of IgA suggests that crosslinking the S protein on the viral surface enhances neutralizing activity either directly or simply through increased apparent affinity. This observation is consistent with the finding that monovalent Fab fragments of serum IgG antibodies are far less potent than the intact antibody (*32*). Whether this effect is due to inter-or intra-spike crosslinking is not known, but it indicates that antibodies or drugs designed to block entry by binding to the RBD could be made more potent by increasing their valency.

A number of different candidate vaccines to SARS-CoV-2 are currently being evaluated in the clinic. Secretory IgA responses may be particularly important to these efforts in that potent dimeric forms of these antibodies are found at the mucosal surfaces where cells are initially targeted by SARS-CoV-2. Thus, even vaccines that elicit modest neutralizing activity in serum may be protective because the secretory polymeric forms of antibodies in mucosa can neutralize the virus. Vaccines that are specifically designed to elicit mucosal IgA responses may be particularly effective preventing SARS-CoV-2 infection (*7*).

## Acknowledgements

We thank all study participants who devoted time to our research; Drs. Barry Coller and Sarah Schlesinger, the Rockefeller University Hospital Clinical Research Support Office and nursing staff; Ivo Lorenz and the Tri-I TDI antibody team for help with BLI measurements. All members of the M.C.N. laboratory for helpful discussions, Maša Jankovic for laboratory support.

## Author contributions

Z.W., J.C.C.L., F.M., S.F. and M.C.N. conceived, designed and analyzed the experiments. Z.W., J.C.C.L., F.M., S.F., C.V., M.C., H.-H.H. L.N. and E.M. carried out all experiments. D.F.R., M. Caskey and C.G. designed clinical protocols. M.C., A.G. and D.O. produced antibodies. T.Y.O., and V.R. performed bioinformatic analysis. C.M.R., T.M. and P.D.B. helped designing the experiments. Z.W., J.C.C.L., F.M., S.F. and M.C.N. wrote the manuscript with input from all co-authors.

## Data and materials availability

Data are provided in table S1, 2, 4. The raw sequencing data associated with Fig. 3 has been deposited at Github (https://github.com/stratust/igpipeline). This study uses data from a database of human shared BCR clonotypes “https://cabrep.c2b2.columbia.edu/home/”, and from ‘cAb-Rep: A Database of Curated Antibody Repertoires for Exploring Antibody Diversity and Predicting Antibody Prevalence’ and ‘High frequency of shared clonotypes in human B cell receptor repertoires’. Computer code to process the antibody sequences are available at GitHub (https://github.com/stratust/igpipeline).

## Materials and Methods

### Human Study participants

Samples were obtained from 149 individuals under a study protocol approved by the Rockefeller University in New York from April 1 through May 8, 2020 as described in (*11*). All participants provided written informed consent before participation in the study and the study was conducted in accordance with Good Clinical Practice and clinical data collection. The study was performed in compliance with all relevant ethical regulations and the protocol was approved by the Institutional Review Board (IRB) of the Rockefeller University.

### Purification and quantification of IgA and IgG from plasma

IgA and IgG were purified from samples with measurable neutralizing activity, against SARS-CoV-2-RBD (*11*). 300µl of plasma was diluted with PBS heat-inactivated (56°C for 1 hr) and incubated with peptide M/Agarose (Invivogen) or Protein G/Agarose (GE lifeSciences) overnight at 4 °C. The suspension was transferred to chromatography columns and washed with 10 column volumes of 1X-PBS. IgA and IgG were then eluted with 1.5ml of 0.1M glycine (pH=3.0) and pH was immediately adjusted to 7.5 with 1M Tris (pH=8.0). 1X-PBS buffer exchange was achieved using Amicon® Ultra centrifugal filters (Merck Millipore) through a 30-kD membrane according to the manufacturer’s instructions. IgA and IgG concentrations were determined by measurement of absorbance at 280nm using a NanoDrop (Thermo Scientific) instrument and samples were stored at 4°C.

### ELISAs

ELISAs to evaluate the IgG or IgA binding to SARS-CoV-2 RBD were performed as previously described using a validated assay (*37, 38*). High binding 96 half well plates (Corning #3690) were coated with 50 µL per well of a 1µg/mL protein solution in PBS overnight at 4?°C. Plates were washed 6 times with washing buffer (1xPBS with 0.05% Tween 20 (Sigma-Aldrich)) and incubated with 170 µL blocking buffer per well (1xPBS with 2% BSA and 0.05% Tween20 (Sigma) for 1 hour at room temperature (RT). Immediately after blocking, monoclonal antibodies or plasma samples were added in PBS and incubated for 1 hr at RT. Plasma samples were assayed at a 1:200 starting dilution and seven additional 3-fold serial dilutions. Monoclonal antibodies were tested at 10 µg/ml starting concentration and 10 additional 4-fold serial dilutions. Plates were washed 6 times with washing buffer and then incubated with anti-human IgG (Jackson Immuno Research 109-036-088) or anti-human IgA (Sigma A0295) secondary antibody conjugated to horseradish peroxidase (HRP) in blocking buffer at 1:5000 or 1:3000 dilution respectively. Plates were developed by addition of the HRP substrate, TMB (ThermoFisher) for 10 minutes (plasma samples) or 4 minutes (monoclonal antibodies), then the developing reaction was stopped by adding 50µl 1M H_2_SO_4_. ODs were measured at 450 nm in a microplate reader (FluoStar Omega, BMG Labtech). For plasma samples, a positive control (plasma from patient COV21, diluted 200-fold in PBS) and negative control historical plasma samples was added in duplicate to every assay plate for validation. The average of its signal was used for normalization of all the other values on the same plate with Excel software prior to calculating the area under the curve using Prism 8 (GraphPad).

### Cell lines

HT1080_Ace2_ cl.14 cells (*27*), 293T_Ace2_ cells (*11*) and VeroE6 kidney epithelial cells were cultured in Dulbecco’s modified Eagle medium (DMEM) supplemented with 10% FCS at 37°C and 5% CO_2_. In addition, medium for Ace2-overexpressing cell lines contained 5 µg/ml blasticidin and medium for VeroE6 cells was supplemented with 1 % nonessential amino acids. All cell lines have been tested negative for contamination with mycoplasma and parental cell lines were obtained from the ATCC.

### Pseudotyped virus neutralization assay

SARS-CoV-2 pseudotyped particles were produced by co-transfection of pSARS-CoV-2 S_trunc_ and pNL4-3ΔEnv-nanoluc in 293T cells (*11, 27*). Four-fold serially diluted purified plasma IgG/IgA from COVID-19 convalescent individuals and healthy donors or monoclonal antibodies were incubated with the SARS-CoV-2 pseudotyped virus for 1 hour at 37?°C. Subsequently, the mixture was incubated with Ace2-expressing cells for 48 hours. HT1080_Ace2_ cl. 14 cells (*27*) were used for plasma-derived IgG/IgA and 293T_Ace2_ cells (*11*) for monoclonal antibodies. Following incubation, cells were washed twice with PBS and lysed with Luciferase Cell Culture Lysis 5x reagent (Promega). Nanoluc Luciferase activity in lysates was measured using the Nano-Glo Luciferase Assay System (Promega) with a GloMax Natigator Microplate Luminometer (Promega). Relative luminescence units obtained were normalized to those derived from cells infected with SARS-CoV-2 pseudotyped virus in the absence of plasma-derived or monoclonal antibodies. The half-maximal and 90% inhibitory concentrations for purified plasma IgG or IgA or monoclonal antibodies (IC50 and IC90) were determined using 4-parameter nonlinear regression (GraphPad Prism).

### Antibody sequencing, cloning and expression

Single B cells were isolated from COV21, COV47 and COV96 patients as previously described(*11*). Briefly, RNA from single cells was reverse-transcribed (SuperScript III Reverse Transcriptase, Invitrogen, 18080-044) using random primers (Invitrogen, 48190011) and followed by nested PCR amplifications and sequencing using the primers for heavy chain that are listed in (table S7) and primers light chains from (*39*). Sequence analysis was performed with MacVector. Antibody cloning from PCR products was performed as previously described by sequencing and ligation-independent cloning into antibody expression vectors (Igγ1-, IGκ-, IGλ-, Igα1 and Igα2) as detailed in (*40*). The Igα1 and Igα2 vectors were from (Invivogen, pfusess-hcha1for IgA1 and pfusess-hcha2m1 for IgA2). J chain plasmid was a gift from Susan Zolla-Pazner. Recombinant monoclonal antibodies were produced and purified as previously described (*39, 41*). Briefly, monoclonal antibodies were produced by transient co-transfection of 293-F cells with human heavy chain and light chain antibody expression plasmids using polyethylenimine (PEI) (Sigma-Aldrich, catalog #408727). Seven days after transfection, supernatants were harvested, clarified by centrifugation and subsequently incubated with Peptide M(Invivogen)/Protein G-coupled sepharose beads (Invivogen, catalog# gel-pdm-5; GE healthcare, 17-0618-05) overnight at 4°C. For dimers, antibodies were produced by transient transfection of Expi293F cells with heavy chain, light chain and J chain expression plasmids at a 1:1:1 ratio. After five days, antibodies were harvested, filtered, incubated with Peptide M overnight and eluted.

### Separation of Dimeric IgA from its Monomeric Form by Size Exclusion Chromatography

A Pre-packed HiLoad™ 16/60 Superdex™ 200 pg (Cytiva, catalog #28989335) on the NGC™ Quest 10 Plus Chromatography System by Bio-Rad was calibrated at room temperature using the HMW Gel Filtration Calibration Kit (Cytiva, catalog #28403842) and IgG. After equilibration of the column with PBS, each concentrated IgA preparation was applied onto the column using a 1 ml-loop at a flow rate of 0.5 ml/min. Dimers of IgA1 or IgA2 were separated from monomers upon an isocratic elution with 70 ml of PBS. The fractions were pooled, concentrated and evaluated by SDS-PAGE using 4 –12% Bis–Tris Novex gels (GenScript catalog #M00652) under reducing and non-reducing conditions followed by a Coomassie blue staining (Expedeon, catalog #ISB1L).

### Microneutralization assay with authentic SARS-CoV-2

Production of SARS-CoV-2 virus was performed as previously described (*11*). This assay was performed as described previously (*11, 42*). VeroE6 cells were seeded at 1×10^4^ cells/well into 96-well plates on the day before infection. IgA monomers and dimers were serially diluted (4-fold) in BA-1, consisting of medium 199 (Lonza, Inc.) supplemented with 1% bovine serum albumin (BSA) and 1x penicillin/streptomycin. The diluted samples were mixed with a constant amount of SARS-CoV-2 and incubated for 1hr at 37°C. The antibody-virus-mix was then directly applied to VeroE6 cells (MOI of ∼0.1 PFU/cell; n=3) and incubated for 22h at 37°C. Cells were subsequently fixed by adding an equal volume of 7% formaldehyde to the wells, followed by permeabilization with 0.1% Triton X-100 for 10 min. After extensive washing, cells were incubated for 1hr at 37°C with blocking solution of 5% goat serum in PBS (catalog no. 005–000-121; Jackson ImmunoResearch). A rabbit polyclonal anti-SARS-CoV-2 nucleocapsid antibody (catalog no. GTX135357; GeneTex) was added to the cells at 1:1,000 dilution in blocking solution and incubated at 4 °C overnight. Goat anti-rabbit AlexaFluor 594 (catalog no. A-11012; Life Technologies) was used as a secondary antibody at a dilution of 1:2,000. Nuclei were stained with Hoechst 33342 (catalog no. 62249; Thermo Scientific) at a 1:1,000 dilution. Images were acquired with a fluorescence microscope and analyzed using ImageXpress Micro XLS (Molecular Devices, Sunnyvale, CA). All experiments involving SARS-CoV-2 were performed in a biosafety level 3 laboratory.

### Biolayer interferometry

BLI assays were performed on the Octet Red instrument (ForteBio) at 30 °C with shaking at 1,000 r.p.m. Epitope binding assays were performed with protein A biosensor (ForteBio 18-5010), following the manufacturer’s protocol “classical sandwich assay”. (1) Sensor check: sensors immersed 30 sec in buffer alone (buffer ForteBio 18-1105). (2) Capture 1st Ab: sensors immersed 10 min with Ab1 at 40 µg/mL. (3) Baseline: sensors immersed 30 sec in buffer alone. (4) Blocking: sensors immersed 5 min with IgG isotype control at 50 µg/mL. (6) Antigen association: sensors immersed 5 min with RBD at 100 µg/mL. (7) Baseline: sensors immersed 30 sec in buffer alone. (8) Association Ab2: sensors immersed 5 min with Ab2 at 40 µg/mL. Curve fitting was performed using the Fortebio Octet Data analysis software (ForteBio).

### Computational analyses of antibody sequences

Antibody sequences were trimmed based on quality and annotated using Igblastn v1.14.0[ref] with IMGT domain delineation system. Annotation was performed systematically using Change-O toolkit v.0.4.5(*43*). Heavy and light chains derived from the same cell were paired, and clonotypes were assigned based on their V and J genes using in-house R and Perl scripts (Fig. 3 A and B). All scripts and the data used to process antibody sequences are publicly available on GitHub (https://github.com/stratust/igpipeline). Nucleotide somatic hypermutation and CDR3 length were determined using in-house R and Perl scripts. For somatic hypermutations, IGHV and IGLV nucleotide sequences were aligned against their closest germlines using Igblastn and the number of differences were considered nucleotide mutations. The average mutations for V genes was calculated by dividing the sum of all nucleotide mutations across all patients by the number of sequences used for the analysis. Hydrophobicity distribution comparisons were calculated as described in (*11*) (Fig. S5). The frequency distributions of human V genes in anti-SARS-CoV-2 antibodies from this study was compared to 131,284,220 IgH and IgL sequences generated by (*44*) and downloaded from cAb-Rep (*45*), a database of human shared BCR clonotypes available at https://cab-rep.c2b2.columbia.edu/. Based on the 81 distinct V genes that make up the 1455 analyzed sequences from Ig repertoire of the three patients present in this study, we selected the IgH and IgL sequences from the database that are partially coded by the same V genes and counted them according to the constant region. The frequencies shown in (Fig. S6) are relative to the source and isotype analyzed. We used the two-sided binomial test to check whether the number of sequences belonging to a specific IgHV or IgLV gene in the repertoire is different according to the frequency of the same IgV gene in the database. Adjusted p-values were calculated using the false discovery rate (FDR) correction. Significant differences are denoted with stars.

---

**Fig. S1.**
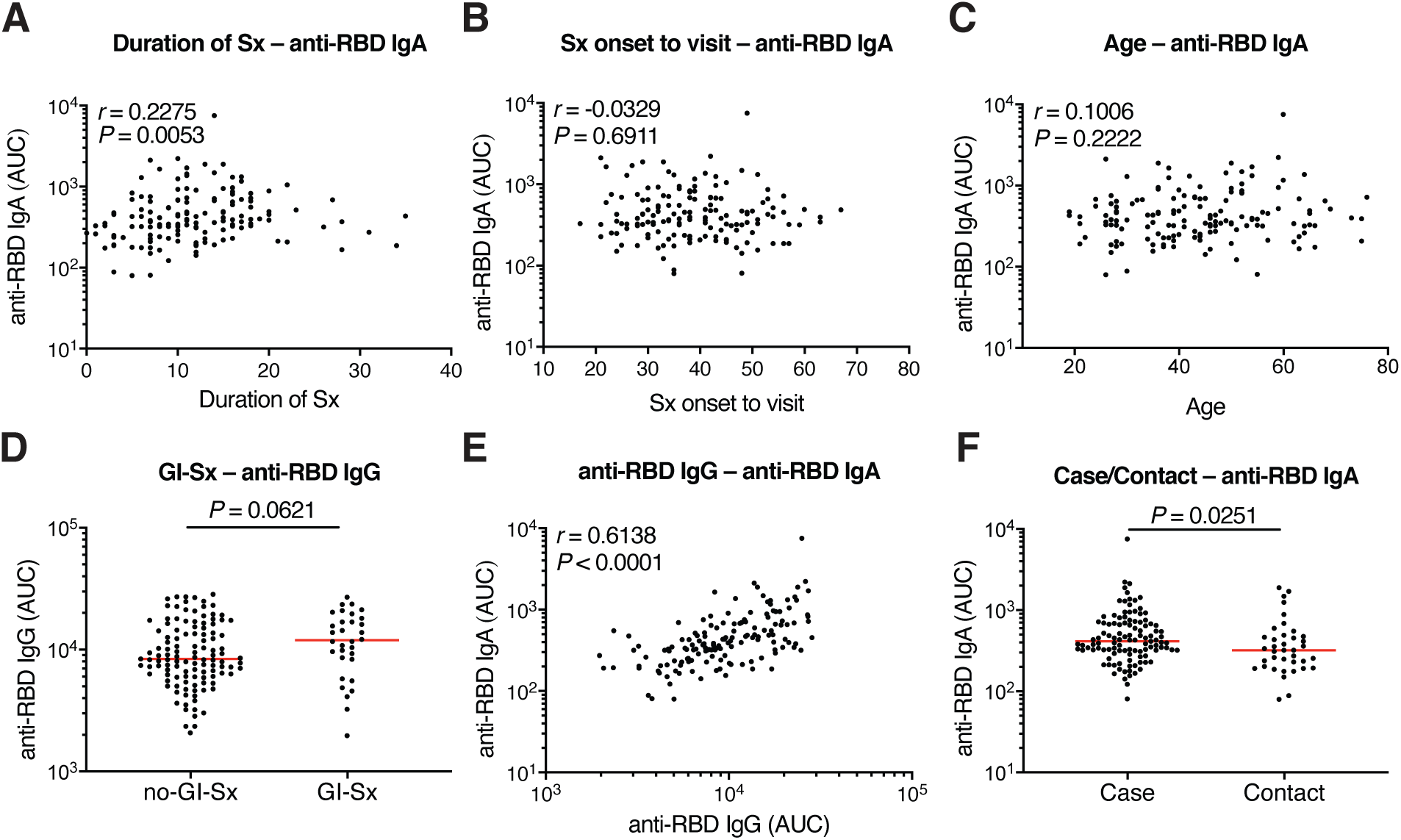
Clinical correlates of plasma IgA antibody titers. (A) Duration of Symptom (Sx) in days (X axis) plotted against normalized AUC for plasma IgA binding to RBD (Y axis). *r* = 0.2275, *P* = 0.0053. (B) Sx onset to time of sample collection in days plotted against normalized AUC for plasma IgA anti-RBD. *r* = −0.0329 and *P* = 0.6911. (C) Age plotted against normalized AUC for plasma IgA anti-RBD. *r* = 0.1006, *P* = 0.2222. (D) Normalized AUC of plasma anti-RBD IgG ELISA for patients with gastrointestinal (GI) symptoms (n=32) and without GI symptoms (n=117); *P* = 0.0621. (E) Normalized AUC of plasma anti-RBD IgG ELISA plotted against normalized AUC for plasma IgA anti-RBD. *r*=0.6138, *P* < 0.0001. (F) Normalized AUC of plasma anti-RBD IgA ELISA for all cases (n = 111) and contacts (n = 38) in the cohort; *P* = 0.0251. For (A-C, E) the correlations were analyzed by two-tailed Spearman’s tests; For (D and F), Horizontal bars indicate median values. Statistical significance was determined using two-tailed Mann–Whitney U-tests.

**Fig. S2.**
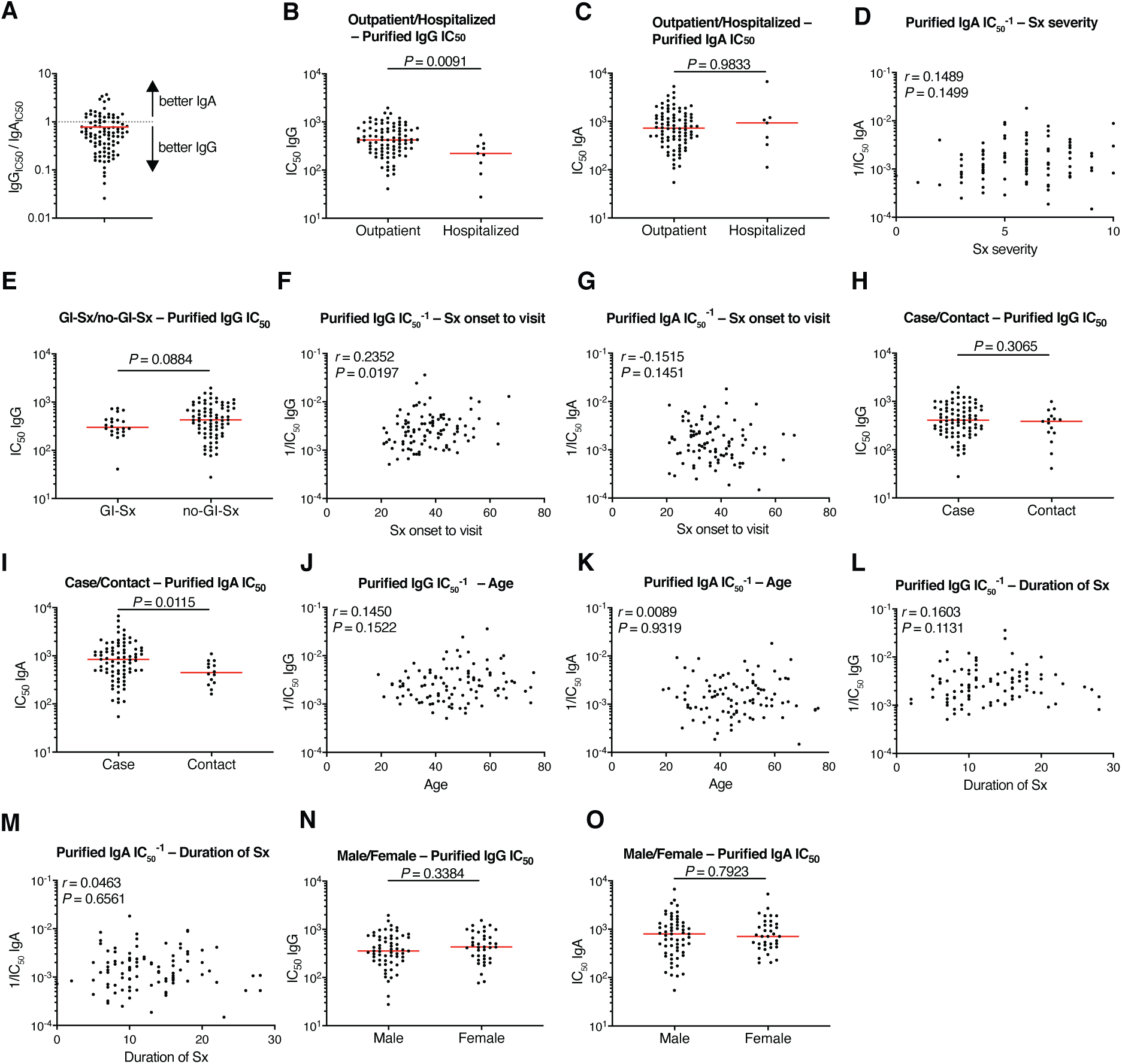
Clinical correlates of plasma IgA/IgG neutralization. (A) Ratio of pseudovirus neutralization IC_50_ values of purified IgG to IgA (n=95). (B, C) Purified plasma IgG (B) and IgA(C) pseudovirus neutralizing IC_50_ values for all outpatient (n = 90) and hospitalized (n = 9) participants in the cohort. (Fig. S2B, *P* = 0.0091) and (Fig. S2C, *P* = 0.9833). (D) Purified plasma IgA 1/IC_50_ values plotted against symptom severity. *r* = 0.1489, *P*=0.1499. (E) Purified plasma IgG IC_50_ values for patients with GI symptoms (n=22) and without GI symptoms (n=77); *p*=0.0884. (F, G) Sx onset to time of sample collection in days plotted against purified plasma IgG (F) and IgA(G) pseudovirus neutralization 1/IC_50_ values. (Fig. S2F, *r* = 0.2352, *P* = 0.0197) and (Fig. S2G, *r* = −0.1515, *P* = 0.1451). (H, I) Purified plasma IgG (H) and IgA(I) pseudovirus neutralization IC_50_ values for all cases (n = 84) and contacts (n = 15) in the cohort. (Fig. S2H, *P* = 0.3065) and (Fig. S2I, *P* = 0.0115). (J, K) Age plotted against purified plasma IgG (J) and IgA(K) pseudovirus neutralization 1/IC_50_ values. (Fig. S2J, *r* = 0.1450, *P* = 0.1522) and (Fig. S2K, *r* = 0.0089, *P* = 0.9319). (L, M) Duration of Symptom (Sx) in days (X axis) plotted against purified plasma IgG (L) and IgA(M) pseudovirus neutralization 1/IC_50_ values. (Fig. S2L, *r* = 0.1603, *P* = 0.1131) and (Fig. S2M, *r* = 0.0463, *P* = 0.6561). (N, O) Purified plasma IgG (N) and IgA(O) pseudovirus neutralization IC_50_ values for males (n=61) and females (n=38). (Fig. S2N, *P*=0.3384) and (Fig.S2O, *P*=0.7923). For (A), horizontal bars indicate mean value. For (B, C, E, H, I, N, O), horizontal bars indicate median values. Statistical significance was determined using two-tailed Mann–Whitney U-tests; For (D, F, G, J-M), the correlations were analyzed by two-tailed Spearman’s tests.

**Fig. S3.**
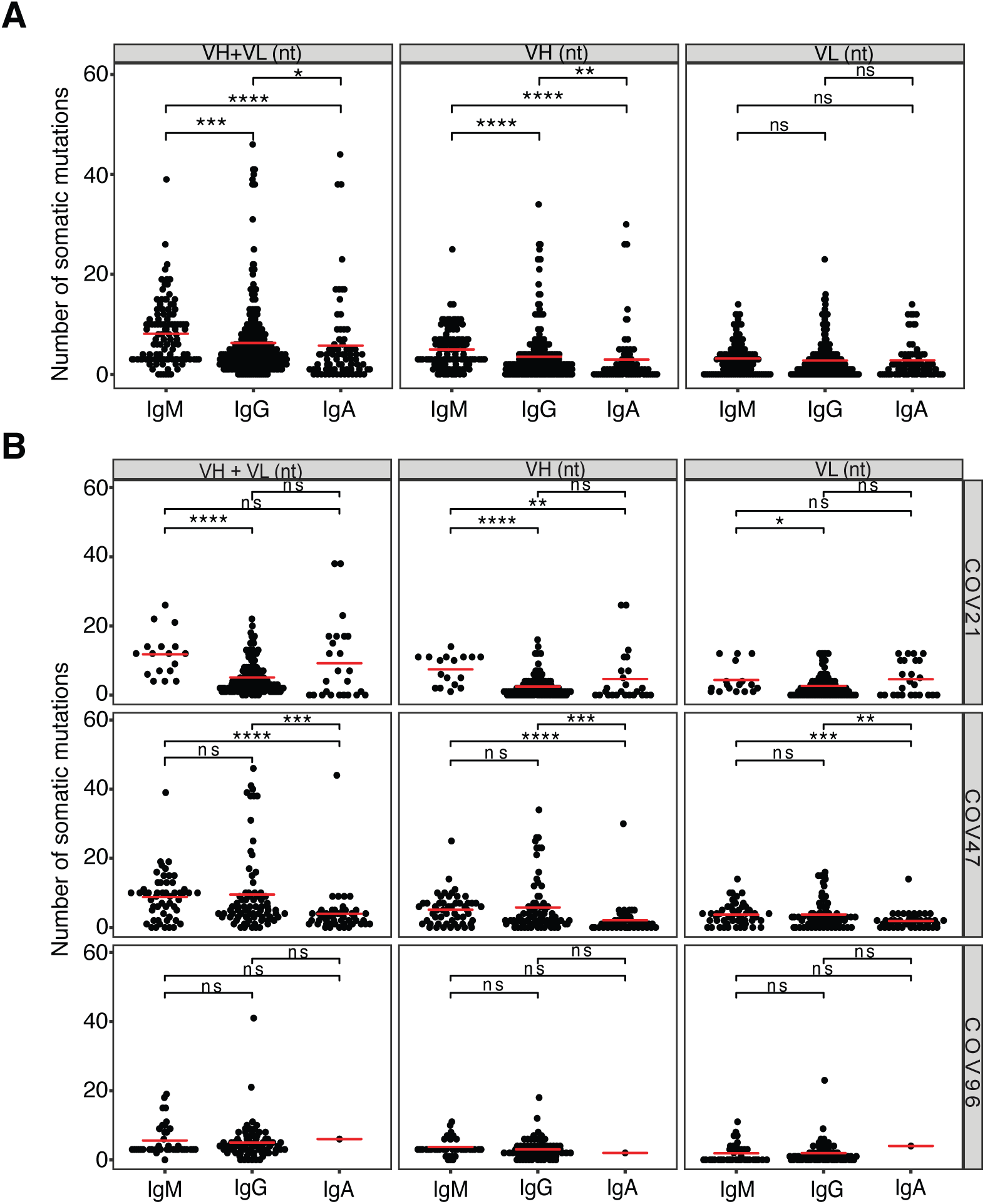
Analysis of antibody somatic hypermutation. (A) The number of somatic nucleotide mutations (Y axis) at the IGVH and IGVL for IgM, IgG and IgA antibodies (X axis), the horizontal bars indicate the mean. The number of antibody sequences was evaluated for both IGVH and IGVL. (n=455). (B) Same as (A) but for each individual.

**Fig. S4.**
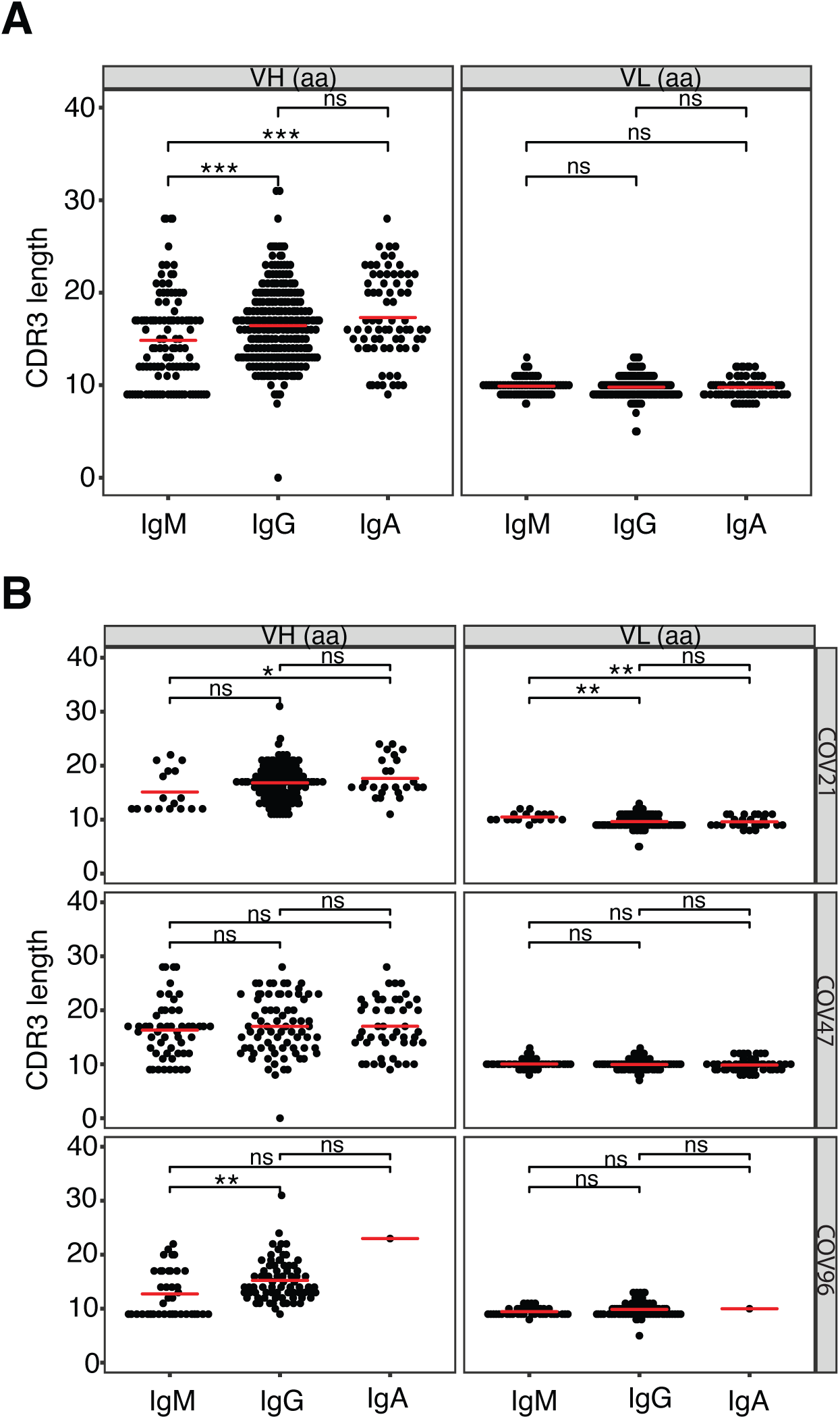
Analysis of antibody CDR3 length. (A) IGVH and IGVL CDR3s length (Y axis) for IgM, IgG and IgA (X axis). (B) Same as (A) but for each individual. The horizontal bars indicate the mean.

**Fig. S5.**
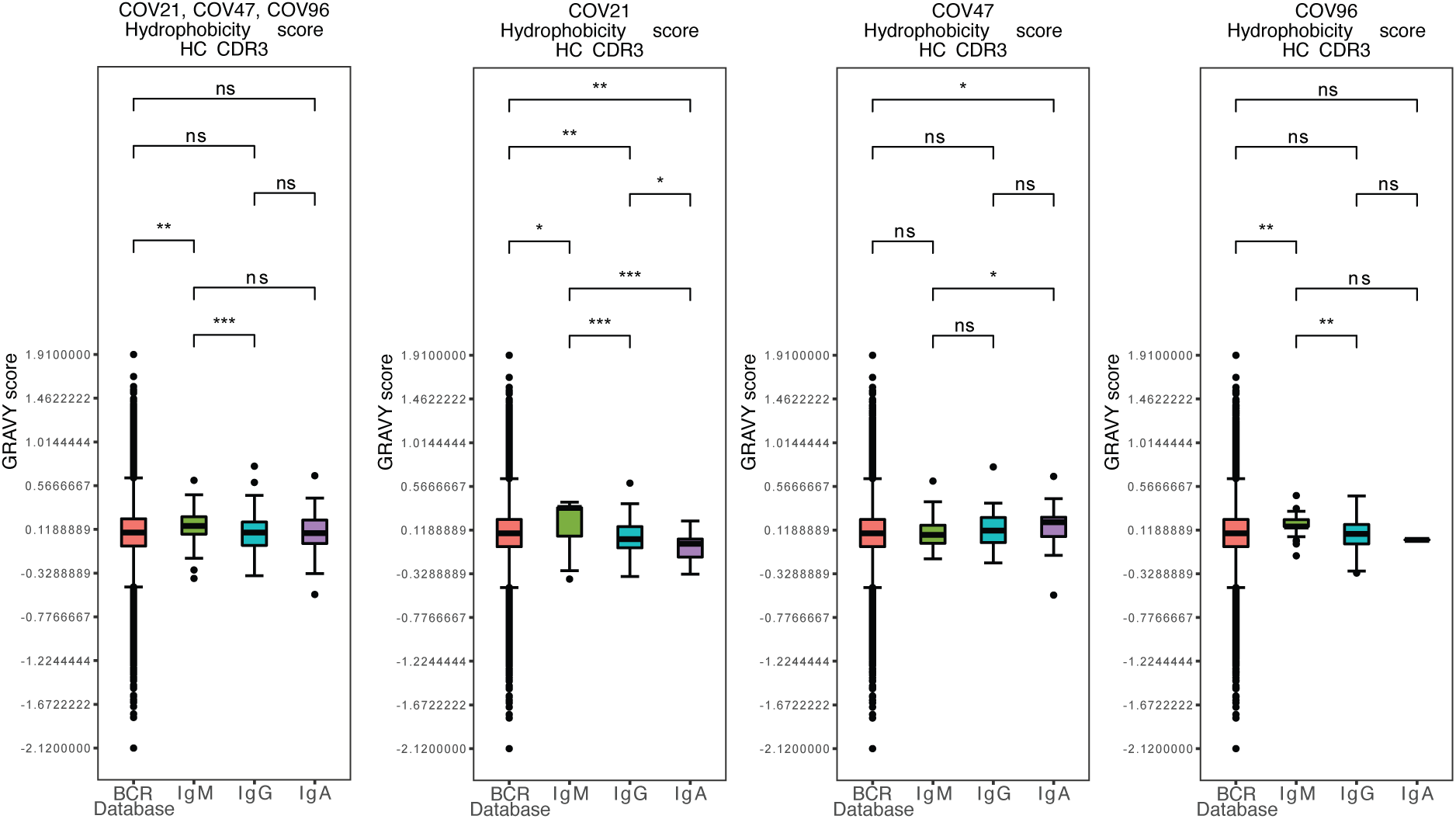
Analysis of antibody CDR3 hydrophobicity. Distribution of the hydrophobicity GRAVY scores at the IGH CDR3 in antibody sequences from this study compared to a public database (see Methods for statistical analysis). The box limits are at the lower and upper quartiles, the center line indicates the median, the whiskers are 1.5x interquartile range and the dots represent outliers.

**Fig. S6.**
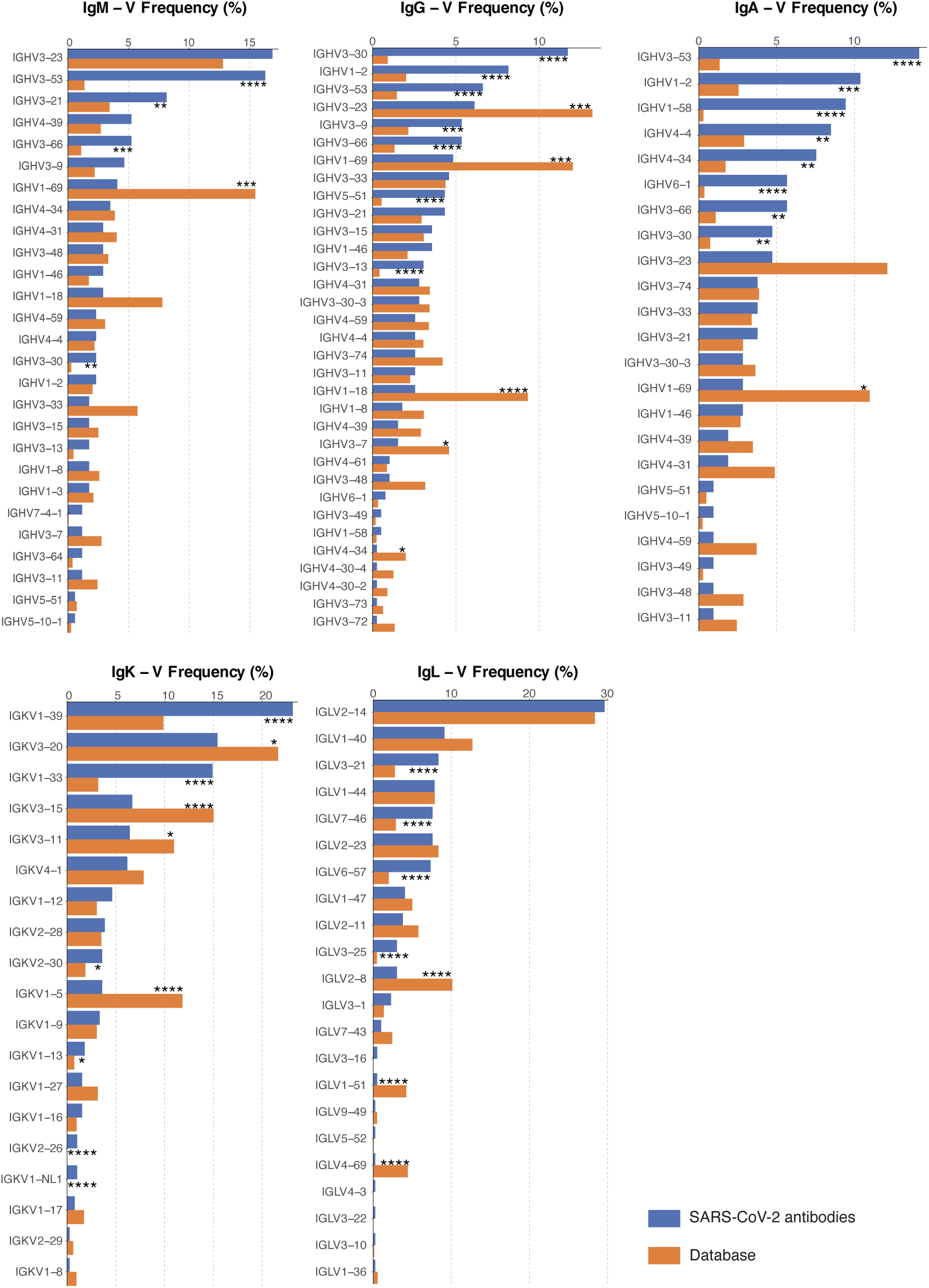
Frequency distributions of human V genes. Comparison of the frequency distributions of human V genes for heavy chain (IgM, IgG and IgA) and light chains of anti-SARS-CoV-2 antibodies from this study and from a database of shared clonotypes of human B cell receptor generated by Cinque Soto et al. (*44*). Statistical significance was determined using the two-sided binomial test. Significant differences are denoted with stars.

**Fig. S7.**
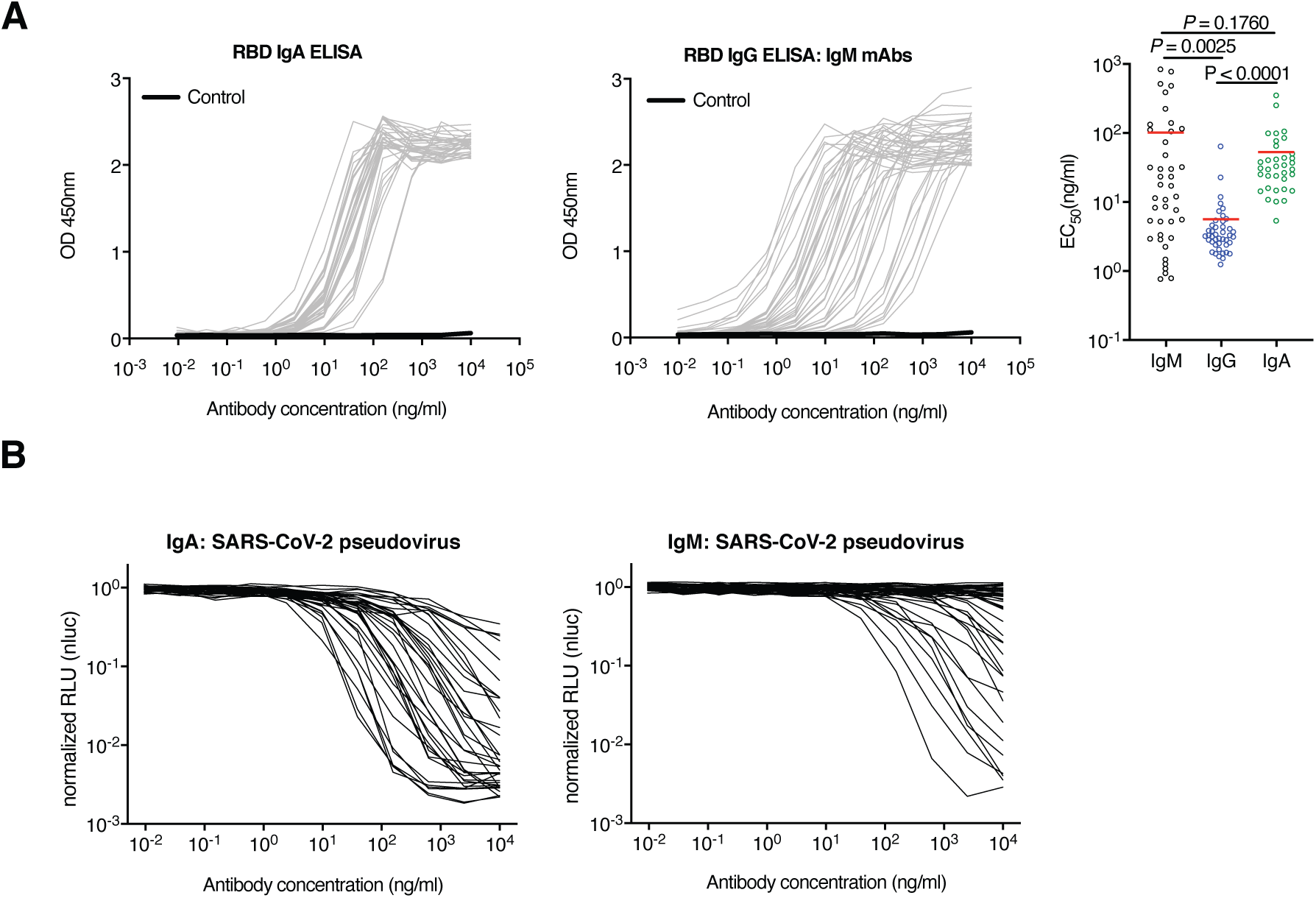
Binding and neutralizing activity of anti-SARS-CoV-2 RBD IgA and IgM monomers. (A) Binding profiles of 35 IgA and 46 IgM monoclonals against RBD. Comparisons of IgM, published IgG (*11*) and IgA EC_50_ values shown as in right panel. Red lines indicate mean value. (B) The normalized relative luminescence values for cell lysates of 293T_ACE2_ cells 48 h after infection with SARS-CoV-2 pseudovirus in the presence of increasing concentrations of monoclonal IgA and IgM antibodies. Statistical analysis was performed using the student’s *t* test.

**Fig. S8.**
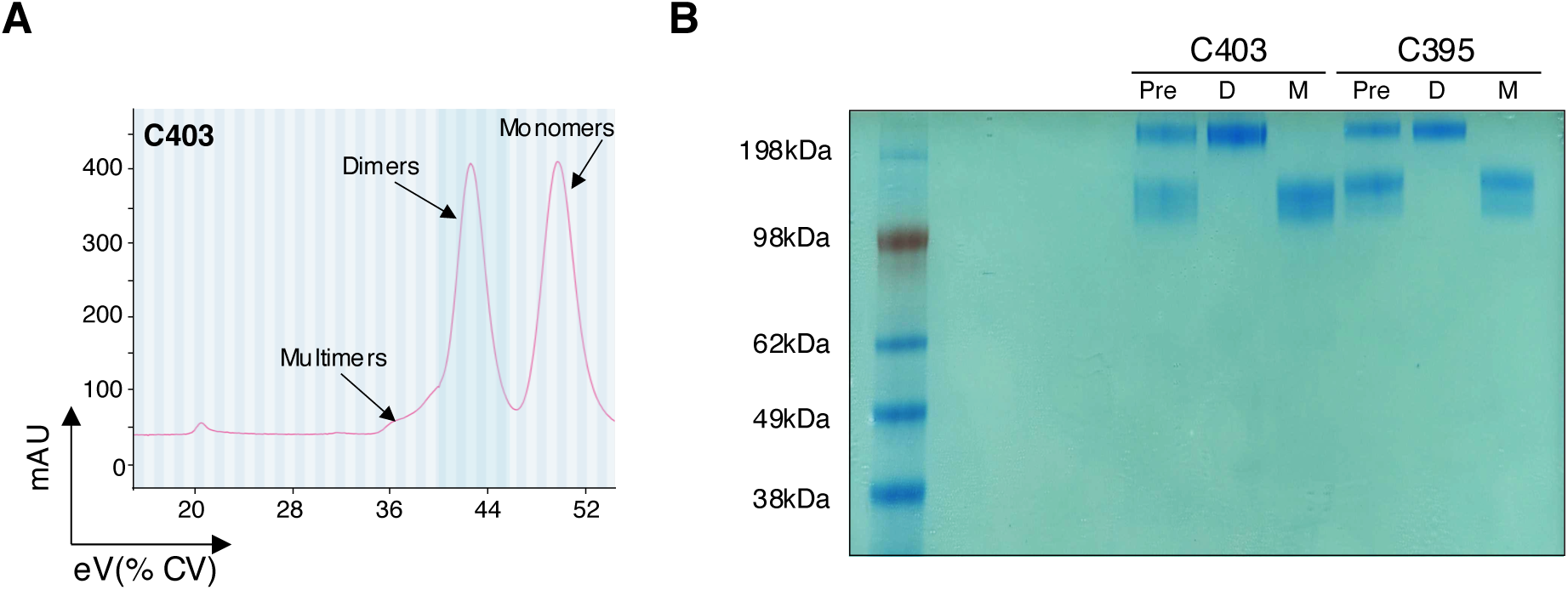
Purification of Dimeric IgA by Size Exclusion Chromatography. (A) Monomers and dimers of IgA1 or IgA2 were separated using a Superdex 200 (Cytiva) with PBS at a flow rate of 0.5 ml/min. Representative example: C403. The X axis is elution volume (eV) as a percent of Column volume. The Y axis is absorption at 280nm (mAU). (B) Coomassie Blue stained non-reducing SDS-PAGE gel of pre-separation antibody mixture (Pre), isolated dimers (D) and monomers (M).

**Fig. S9.**
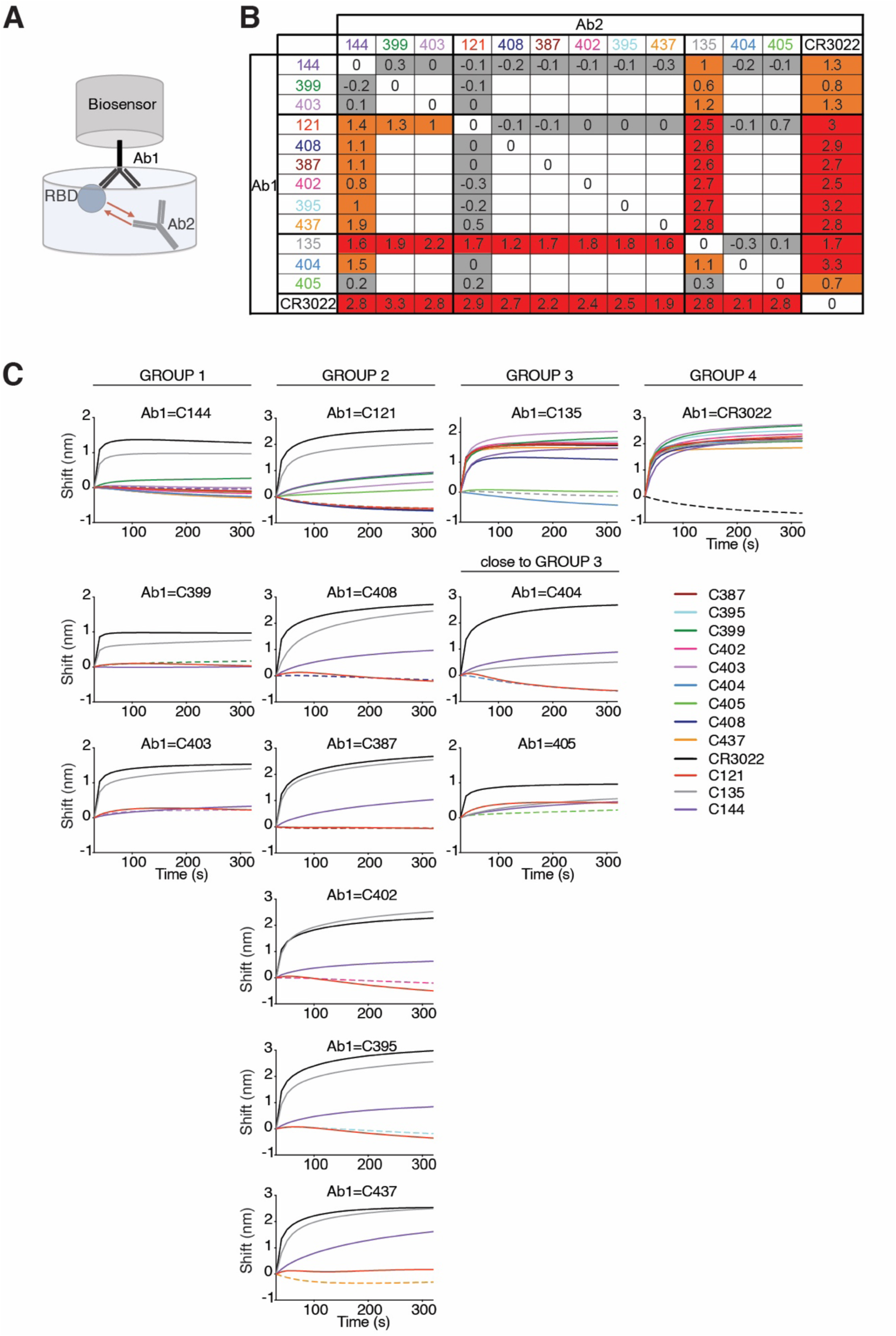
Biolayer interferometry experiment. (A) Diagrammatic representation of biolayer interferometry experiment. (B) The table displays the shift in nanometers after second antibody (Ab2) binding to the antigen in the presence of the first antibody (Ab1). Values are normalized by the subtraction of the autologous antibody control. (C) Second antibody (Ab2) binding to preformed first antibody (Ab1)–RBD complexes. Dotted line denotes when Ab1 and Ab2 are the same, and Ab2 is according to the colour-coding in Fig. 4B (right panel).

**Fig. S10.**
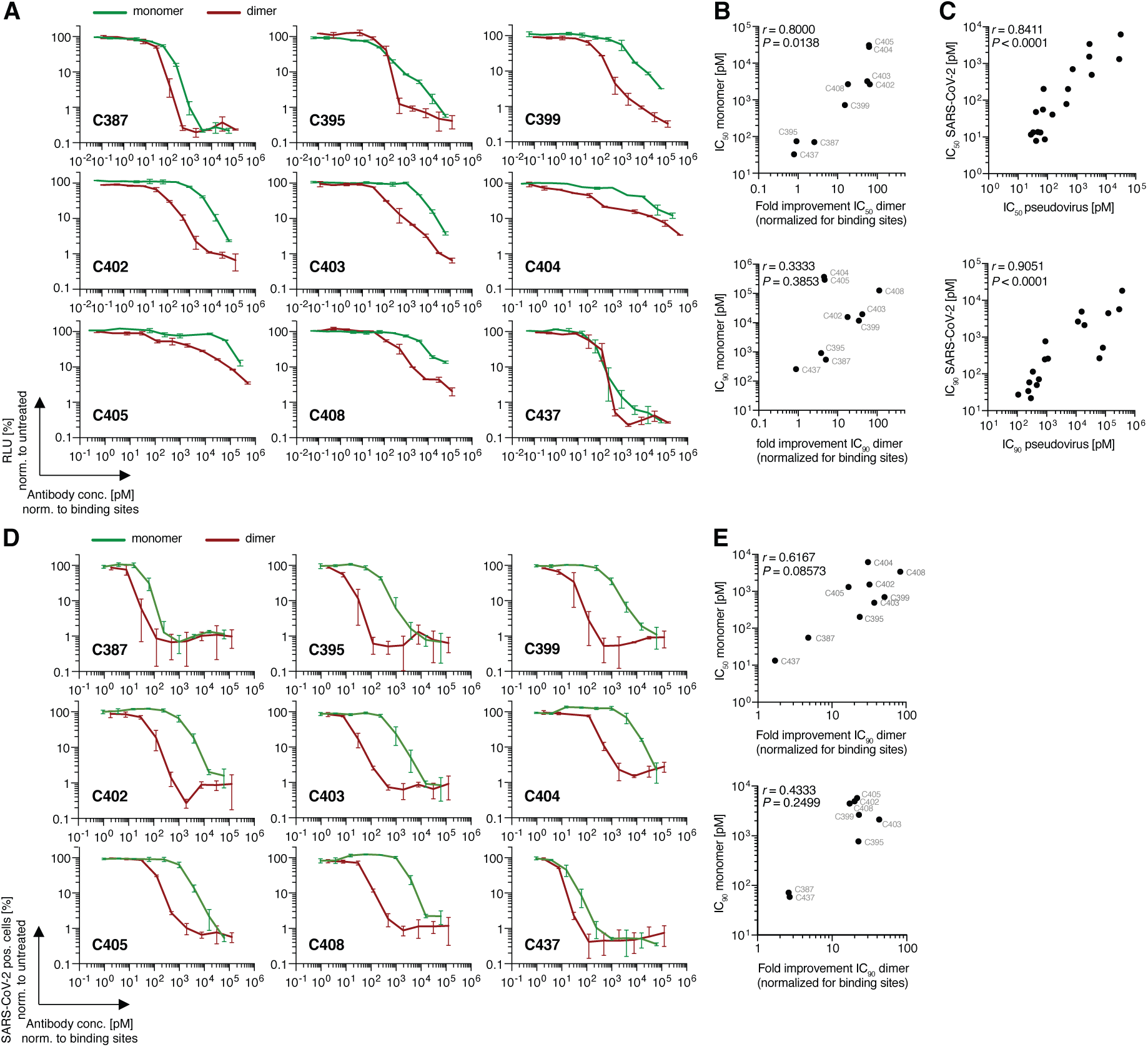
Neutralizing activity of monoclonal monomeric and dimeric IgAs. (A) The normalized relative luminescence values for cell lysates of 293T_ACE2_ cells 48 h after infection with SARS-CoV-2 pseudovirus in the presence of increasing concentrations of monoclonal antibodies C387, C395, C399, C402, C403, C404, C405, C408, C437 in their monomeric (green curves) and dimeric (red curves) form. (B) Fold improvement of the IC_50_ (upper panel) and IC_90_ (lower panel) values of dimeric IgA to monomeric IgA (X axis) plotted against IC_50_ (*r* = 0.8000, *P* = 0.0138), IC_90_ (*r* = 0.3333, *P* = 0.3853) values of monomeric IgAs. (C) IC_50_ (upper panel) and IC_90_ (lower panel) values of dimeric and monomeric IgAs determined by pseudovirus neutralization assay (x axis) plotted against IC_50_ (*r* = 0.8411, *P* < 0.0001) and IC_90_ (*r* = 0.9051, *P* < 0.0001) values determined by authentic SARS-CoV-2 neutralization assay (y axis). (D) SARS-CoV-2 neutralization assay. The normalized percentage of SARS-CoV-2 positive VeroE6 cells 48 h after infection with SARS-CoV-2 authentic virus in the presence of increasing concentrations of abovementioned antibodies in their dimeric and monomeric form. (E) Fold improvement of the IC_50_ (upper panel) and IC_90_ (lower panel) values of dimeric IgA to monomeric IgA (X axis) plotted against IC_50_ (*r* = 0.6167, *P* = 0.08573), IC_90_ (*r* = 0.4333, *P* = 0.2499) values of monomeric IgAs. Correlations were analyzed by two-tailed Spearman’s tests.

**Table S1. Sequences of anti-SARS-CoV-2 antibodies**

Auxiliary Supplementary Material.

**Table S2. Sequences of antibodies from isotype shared clones**

Auxiliary Supplementary Material.

**Table S3.** Inhibitory concentrations of monoclonal antibodies from isotype shared clones

| Patient ID | IgM | IC50(ng/ml) | IC90(ng/ml) | IgG | IC50(ng/ml)* | IC90(ng/ml)* | IgA | IC50(ng/ml) | IC90(ng/ml) |
| --- | --- | --- | --- | --- | --- | --- | --- | --- | --- |
| COV21 | - |  |  | CG002 | 8.88 | 37.61 | CA386 | 5.76 | 123.33 |
|  | - |  |  | CG005 | 60.49 | 205.20 | CA387 | 9.68 | 129.87 |
| COV47 | - |  |  | CG144 | 6.91 | 29.66 | CA394 | 13.06 | 371.86 |
|  | CM169 | UD | UD | CG148 | >1000 | >1000 | - |  |  |
|  | CM170 | 5806 | 37082 | CG171 | 5250 | 17156 | CA457 | 1721.6 | 298325.6 |
|  | - |  |  | CG379 | 126.98 | 2368.18 | CA403 | 23.88 | 126.05 |
|  | CM381 | UD | UD | CG160 | >1000 | >1000 | - |  |  |
|  | CM349 | 844.59 | 26446.73 | CG380 | 2.94 | 35.96 | - |  |  |
|  | CM311 | 126.85 | 846.13 | CG151 | 31.79 | >1000 | CA390 | 417.42 | 46597.44 |
| COV96 | CM194 | UD | UD | CG382 | 42.92 | 122.33 | - |  |  |
|  | CM365 | 1226.09 | 8268.46 | CG202 | >1000 | >1000 | - |  |  |

UD=Undetectable
\*(Robbiani et al. 2020)

**Table S4. Sequences of cloned recombinant antibodies**

Auxiliary Supplementary Material.

**Table S5. Effective and inhibitory concentrations of monoclonal antibodies**

Auxiliary Supplementary Material.

**Table S6.** Inhibitory concentrations of monoclonal IgA monomers and dimers

| Antibody ID | SARS-CoV-2 pseudovirus |  |  |  | SARS-CoV-2 |  |  |  |
| --- | --- | --- | --- | --- | --- | --- | --- | --- |
|  | IC50 (pM) |  | IC90 (pM) |  | IC50 (pM) |  | IC90 (pM) |  |
|  | monomer | dimer | monomer | dimer | monomer | dimer | monomer | dimer |
| C387 | 70.68 | 27.46 | 543.91 | 108.06 | 55.74 | 11.64 | 71.46 | 27.37 |
| C395 | 74.15 | 81.64 | 909.67 | 239.03 | 203.72 | 8.59 | 769.49 | 34.10 |
| C399 | 722.84 | 47.48 | 11692.05 | 339.46 | 700.38 | 13.67 | 2636.98 | 114.74 |
| C402 | 2652.52 | 40.86 | 15603.86 | 874.99 | 1536.24 | 47.90 | 4939.53 | 247.79 |
| C403 | 3222.29 | 57.62 | 19499.80 | 461.45 | 491.11 | 13.18 | 2115.67 | 49.85 |
| C404 | 31112.25 | 502.11 | 371134.20 | 82318.65 | 6182.08 | 201.78 | 18271.60 | 514.05 |
| C405 | 27801.09 | 444.36 | 294918.48 | 62867.98 | 1312.88 | 78.57 | 5725.29 | 266.04 |
| C408 | 2691.09 | 147.89 | 126130.74 | 1114.44 | 3392.62 | 40.51 | 4458.13 | 259.84 |
| C437 | 32.67 | 41.18 | 258.08 | 292.35 | 13.32 | 7.84 | 58.85 | 21.82 |

*IC50/90 values for dimers were adjusted for number of binding sites*

**Table S7. Primers**

Auxiliary Supplementary Material.

